# Rational engineering of an erythropoietin fusion protein to treat hypoxia

**DOI:** 10.1101/2021.06.24.449789

**Authors:** Jungmin Lee, Andyna Vernet, Nathalie G. Gruber, Kasia M. Kready, Devin R. Burrill, Jeffrey C. Way, Pamela A. Silver

## Abstract

Erythropoietin enhances oxygen delivery and reduces hypoxia-induced cell death, but its pro-thrombotic activity is problematic for use of erythropoietin in treating hypoxia. We constructed a fusion protein that stimulates red blood cell production and neuroprotection without triggering platelet production, a marker for thrombosis. The protein consists of an anti-glycophorin A nanobody and an erythropoietin mutant (L108A). The mutation reduces activation of erythropoietin receptor homodimers that induce erythropoiesis and thrombosis, but maintains the tissue-protective signaling. The binding of the nanobody element to glycophorin A rescues homodimeric erythropoietin receptor activation on red blood cell precursors. In a cell proliferation assay, the fusion protein is active at 10^−14^ M, allowing an estimate of the number of receptor–ligand complexes needed for signaling. This fusion protein stimulates erythroid cell proliferation *in vitro* and in mice, and shows neuroprotective activity *in vitro*. Our erythropoietin fusion protein presents a novel molecule for treating hypoxia.

## Introduction

Erythropoietin (EPO) stimulates red blood cell (RBC) production in response to hypoxia. It inhibits apoptosis of late-stage erythroid precursors (e.g. CFU-E, BFU-E) and promotes their proliferation and maturation into the fully committed erythroid lineage. Healthy human adult kidneys constitutively produce EPO at low levels, maintaining ∼1–5 pM of circulating EPO under normoxic conditions to sustain constant hemoglobin levels (Elliott *et al*., 2014). In response to hypoxic stress or massive blood loss, EPO production is stimulated and the number of circulating erythrocytes increases, allowing for more efficient tissue oxygenation (Ghezzi and Brines, 2004).

EPO, like other cytokines and hormones, is pleiotropic and performs several other biological functions in addition to hematopoiesis. Functional EPO receptors (EPORs) are expressed in many tissues other than erythroid precursors, such as endothelial cells, cardiomyocytes, and cells of the central nervous system (Masuda *et al*., 1994; Ogunshola and Bogdanova, 2013; Hernandez *et al*., 2017). Deletion of EPORs in mouse embryos results not only in impaired erythropoiesis, but also in developmental defects in the heart, the vasculature, and the brain (Ogunshola and Bogdanova, 2013). Existence of functional EPORs in non-hematopoietic tissues suggests that EPO activates EPORs in different contexts to induce biological activities that are independent of erythropoiesis.

Non-hematopoietic functions of EPO include enhancement of blood clotting and tissue protection in response to hypoxia. These functions suggest that EPO mediates the body’s response to hemorrhage, rather than simply being an RBC-producing hormone. When an animal is wounded, the immediate response by the body should be to stop bleeding, increase RBC production, promote tissue oxygenation and ensure tissue survival until oxygen levels return to baseline. Pro-thrombotic effects have been observed as adverse side effects of EPO in the treatment of anemia. Chronic kidney failure patients receiving EPO exhibit higher incidences of strokes, hypertension and death (Drueke *et al*., 2006; Singh *et al*., 2006; Pfeffer *et al*., 2009). Cancer patients treated with EPO had accelerated tumor growth and lower survival rate, possibly due to EPORs on cancer cells themselves, increased tumor angiogenesis, and deep vein thrombosis (Henke *et al*., 2003; Okazaki *et al*., 2008; Yasuda *et al*., 2003). EPO’s tissue-protective effects in response to hypoxia have also been shown in animal models and are suggested in several clinical studies (Ehrenreich *et al*., 2002; Ehrenreich *et al*., 2007; Aloizos *et al*., 2015). Intravenous injections of high doses of EPO significantly reduced infarct size and serum markers of brain damage in acute ischemic stroke patients (Ehrenreich *et al*., 2002), and improved motor and cognitive function in multiple sclerosis patients (Ehrenreich *et al*., 2007). EPO treatment also resulted in a lower mortality rate and improved neurological recovery amongst traumatic brain injury (TBI) patients (Aloizos *et al*., 2015). The protective activity of EPO is general to all cellular insults tested so far, including hypoxia, TBI and neuronal excitotoxicity (Fantacci *et al*., 2006; Robinson *et al*., 2018; Park *et al*., 2011).

Due to its erythropoietic and tissue-protective functions, EPO holds great promise as a therapeutic for various conditions that cause hypoxia, such as chronic obstructive pulmonary disease (COPD), right-side heart failure and viral infection that requires use of a ventilator. However, two major challenges have limited the clinical use of EPO for tissue protection resulting from hypoxia. First, EPO has a pro-thrombotic effect that is observed at low doses, while the tissue-protective effect requires much higher doses. Thus, doses at which EPO might be effective for tissue protection are considered unsafe. Second, EPO (30.4 kDa) has a short plasma half-life of ∼8 hours after a single intravenous injection in humans (Bunn, 2013). Its poor pharmacokinetic profile necessitates frequent dosing to maintain the high levels of EPO required for efficacy.

EPO acts through two distinct receptor complexes (Fig. 1A and B). RBC production and clotting is mediated via EPOR homodimers, whereas the angiogenic and tissue-protective activities of EPO are thought to be regulated by heterodimers of EPOR and the co-receptor CD131 (also known as the receptor common beta subunit) (Hanazono *et al*., 1995; Brines *et al*., 2004; Leist *et al*., 2004; Bennis *et al*., 2012). EPO monomers bind to EPOR homodimers through a strong interaction (K_D_ = 1 nM) on one face involving residues such as N147 and R150 (the ‘strong face’) (Fig. 1A, C and D), and through a weak interaction (K_D_ = 1 μM) on another face involving residues such as S100, R103, S104 and L108 (the ‘weak face’) (Fig. 1A, C and E) (Elliott *et al*., 1997; Syed *et al*., 1998). Tissue-protective signaling through putative EPOR–CD131 heterodimers is thought to involve EPO binding to EPOR through its strong face and an interaction through CD131 that is not well understood (Fig. 1B). This configuration is inferred by the fact that while weak-face mutations (e.g. S100E and R103E) disrupt EPOR homodimer signaling (Leist *et al*., 2004; Elliott *et al*., 1997) and RBC production, there is essentially no effect on neuroprotective signaling (Gan *et al*., 2012). Specifically, Gan *et al*. (2012) introduced nine mutations on the weak face of EPO, and found that all such mutant proteins mediated neuroprotection – i.e. none of the mutations disrupted a possible interaction with CD131. Thus, it appears that the weak face of EPO can be arbitrarily manipulated for protein engineering purposes and still maintain its tissue-protective function.

**Fig. 1.**
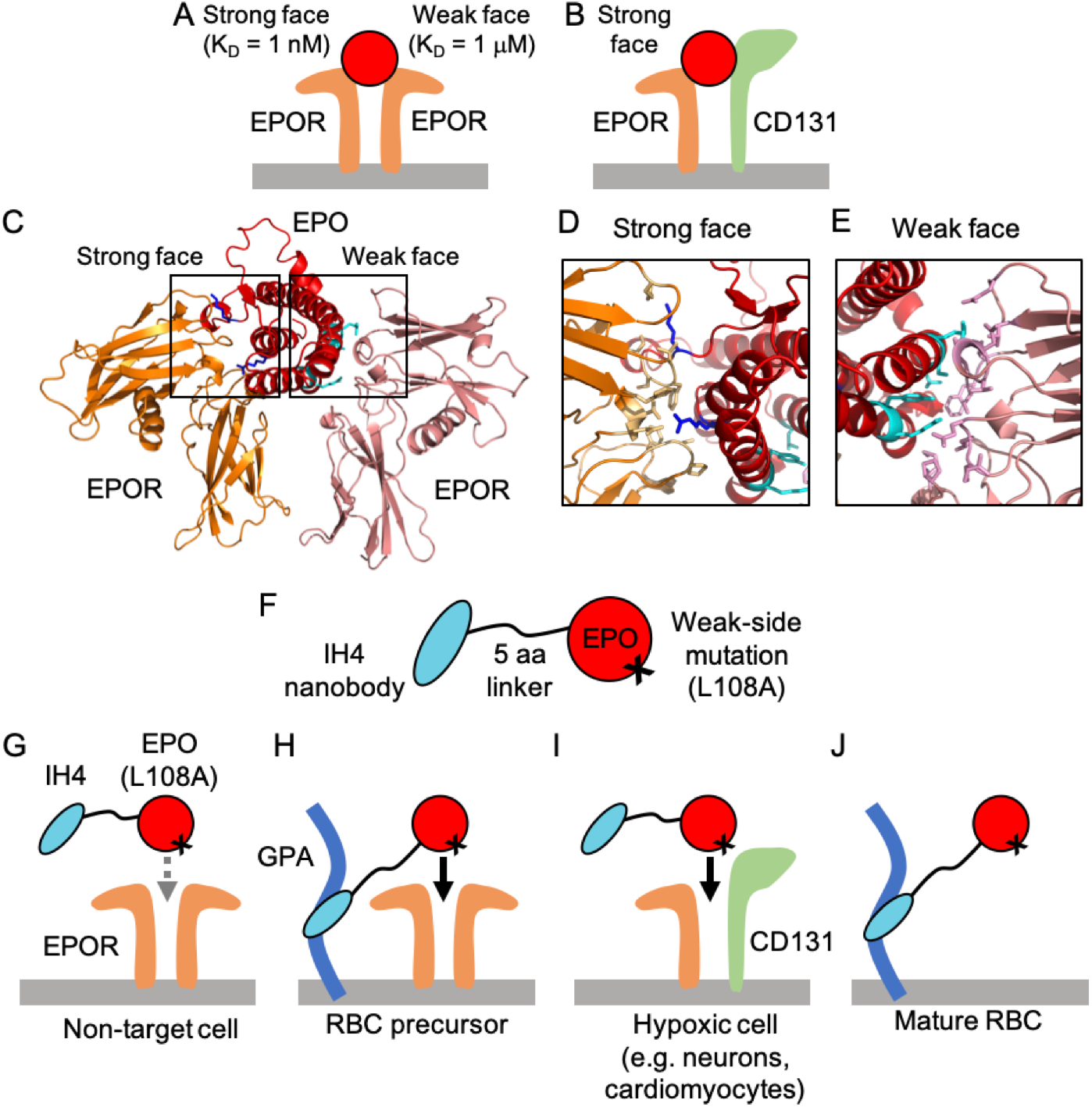
Design rationale for EPO-H fusion protein. (**A–E**) Protein interactions of natural EPO with homodimeric EPOR and heterodimeric EPOR–CD131. **(A)** EPO binds asymmetrically to homodimeric EPOR via two distinct binding interfaces: the strong face (K_D_ = 1 nM) and the weak face (K_D_ = 1 μM). **(B)** EPO can also bind to EPOR–CD131 receptors via its strong face. **(C)** Protein structure of EPO interacting with homodimeric EPOR (PDB ID: 1EER). Zoom-in of the **(D)** strong and (**E)** weak binding interfaces. For receptor binding and activity, critical EPO residues are shown in **(D)** blue sticks (top: K45, bottom: R150) and **(E)** cyan sticks (top to bottom: R103, S104, L108, Y15, R14). EPOR residues that are within 4 Å of these residues are shown in light yellow and pink sticks in **(D)** and **(E)**, respectively. **(F)** The EPO-H fusion protein consists of the IH4 nanobody, which targets GPA-expressing cells, attached to a mutant EPO by a five-amino acid linker. **(G)** Mutant EPO(L108A) has weakened affinity for homodimeric EPOR, and thus, has little effect on non-target cells that lack GPA. **(H)** On erythropoietic target cells that express both GPA and EPOR, the binding of IH4 to GPA localizes the fusion protein to the target cell surface and allows activation of homodimeric EPOR. **(I)** The L108A mutation in the EPO element does not disrupt EPO interaction with CD131. As a result, IH4-EPO(L108A) can induce tissue-protective activity via a heterodimeric EPOR–CD131 receptor complex. **(J)** IH4-EPO(L108A) also binds to mature RBCs via GPA, thereby extending its plasma half-life.

We previously constructed ‘chimeric activator’ proteins in which a mutated EPO with lower receptor affinity is fused to an antibody element that binds to glycophorin A (GPA) (Taylor *et al*., 2010; Burrill *et al*., 2016; Lee *et al*., 2020). Burrill *et al*. (2016) demonstrated that a weakened form of EPO with a mutation in the strong face (R150A) that is also fused to an anti-GPA antibody element can specifically activate production of RBCs and not platelets. Lee *et al*. (2020) demonstrated that such an anti-GPA/EPO(mutant) fusion protein can specifically activate RBC formation without stimulation of blood clotting, provided that the fusion protein cannot mediate adhesion of cells bearing GPA (e.g. RBCs) and other cells bearing EPORs. The results of Lee *et al*. (2020) also showed a correlation between stimulation of platelet production and stimulation of thrombosis, indicating that enhancement of platelet formation could be used as a surrogate marker for EPO-induced thrombosis in these studies. The mutations used in those studies affected the strong face of EPO and those fusion proteins are therefore expected to affect formation of EPOR homodimers and EPOR–CD131 heterodimers. These engineered molecules stimulate RBC formation without activating thrombosis or tissue-protective activity. The present work describes the design of a new molecule that is able to stimulate both RBC production and tissue protection without stimulating platelet formation, a surrogate marker of the pro-thrombotic side effect of EPO.

## Results

### Rational design of EPO fusion proteins to address hypoxia

Our work aims to improve the pharmacokinetics and therapeutic window of EPO, so to harness both its erythropoietic and tissue-protective effects while avoiding thrombosis. To achieve this goal, we designed EPO fusion proteins (EPO-H; H for hypoxia) based on the concept of a ‘chimeric activator’ previously developed (Taylor *et al*., 2010; Burrill *et al*., 2016; Lee *et al*., 2020).

EPO-H consists of the nanobody element IH4, which binds to the target antigen GPA, and a mutated version of EPO, fused via a flexible five-amino acid linker (Fig. 1F). We hypothesized that a mutation in the EPO element could weaken its affinity to homodimeric EPOR, thereby avoiding undesired pro-thrombotic effects triggered by homodimeric EPOR signaling on non-target cells (Fig. 1G). The desired erythropoietic activity is rescued by targeted EPO activity on RBC precursor cells directed by the binding of the antibody element, IH4, to the target antigen, GPA (Fig. 1H). This way, EPO activates homodimeric EPORs only on RBC precursors, mitigating the unwanted thrombotic side effects via non-target cells.

At the same time, it is important to ensure that the same mutation in the EPO element does not disrupt EPO binding to heterodimers of EPOR and the co-receptor CD131 when the tissue-protective activity is desired (Fig. 1I). Several EPO mutants were designed based on previous mutagenesis studies (Elliott *et al*., 1997; Gan *et al*., 2012). The predicted behaviors of these EPO mutants, either alone or when fused to an anti-GPA antibody element, are outlined in Table I. As part of the design strategy, we use “leaky mutations” that reduce but do not abolish binding; in practice these are mutations in which an amino acid with a long side chain is replaced by one with a shorter side chain, so that no steric hindrance results and binding is possible. As controls, we also use “tight mutations” in which a side chain is lengthened or the charge of a side chain is reversed, so that a binding activity would be completely lost due to the specific mutation. Because the binding mode of EPO to EPOR–CD131 is not elucidated, unlike that of EPO to homodimeric EPOR, several single point mutations were made in both of the known EPOR contact regions (strong and weak faces, each with K_D_ of 1 nM and 1 μM, respectively). Two residues on the strong face, K45 and R150, and five residues on the weak face, R14, Y15, R103, S104 and L108, were mutated and tested for targeted erythropoietic and tissue-protective activities (Table II and Fig. S1).

**Table I.** Table depicting predicted properties of wildtype EPO and engineered EPO-H variants. Predicted properties: RBC production, thrombosis, tissue protection, and expected half-life. “*+*” = increase, “*–*” = decrease or no effect.

|  | Mutation | Protein | RBC production | Thrombosis | Tissue protection | Expected Half-life |
| --- | --- | --- | --- | --- | --- | --- |
|  | Wildtype | EPO | + | + | + | Short |
| Strong-face mutation | R150A (Leaky) | EPO(R150A) | – | – | – | Short |
|  |  | IH4-EPO(R150A) | + | – | – | Extended |
|  | K45D (Tight) | EPO(K45D) | – | – | – | Short |
|  |  | IH4-EPO(K45D) | – | – | – | Extended |
| Weak-face mutation | L108A (Leaky) | EPO(L108A) | – | – | + | Short |
|  |  | IH4-EPO(L108A) | + | – | + | Extended |
|  | S104I (Tight) | EPO(S104I) | – | – | + | Short |
|  |  | IH4-EPO(S104I) | – | – | + | Extended |

**Table II.** *In vitro* stimulation of TF-1 cell proliferation by EPO mutants and their fusion to IH4. *N* indicates the number of repeat experiments, each containing three replicates. N.D. = Not determined. N.A. = Not active.

|  | Protein | EPO |  |  | IH4-EPO |  |  |
| --- | --- | --- | --- | --- | --- | --- | --- |
| | | *N* | Log($EC_{50}$) (M)<br>± S.D. | $EC_{50}$ relative<br>to epoetin alfa | *N* | Log($EC_{50}$) (M)<br>± S.D. | $EC_{50}$ relative<br>to epoetin alfa |
| Control | Epoetin alfa (Epogen®) | 21 | -10.22 ± 0.32 | 1 | 0 | N.D. | N.D. |
|  | Darbepoetin (Aranesp®) | 10 | -9.06 ± 0.27 | 14.20 | 0 | N.D. | N.D. |
| Strong Face | K45D | 4 | -6.68 ± 0.65 | 3425.96 | 2 | -9.21 ± 0.49 | 10.04 |
|  | R150A | 8 | -8.14 ± 0.06 | 119.32 | 9 | -10.41 ± 0.29 | 0.64 |
| Weak Face | S104I | 2 | N.A. | N.A. | 2 | N.D. | N.D. |
|  | R14E | 1 | N.A. | N.A. | 3 | N.A. | N.A. |
|  | R14Q | 1 | N.A. | N.A. | 2 | N.A. | N.A. |
|  | R14N | 1 | N.A. | N.A. | 2 | N.A. | N.A. |
|  | Y15I | 2 | N.A. | N.A. | 2 | N.A. | N.A. |
|  | R103I | 1 | N.A. | N.A. | 3 | N.A. | N.A. |
|  | R103Q | 2 | N.A. | N.A. | 3 | N.A. | N.A. |
|  | R103K | 6 | -9.93 ± 0.40 | 1.94 | 8 | -13.16 ± 1.74 | 1.14x10<sup>-3</sup> |
|  | L108A | 5 | N.A. | N.A. | 5 | -13.74 ± 1.54 | 2.99x10<sup>-4</sup> |

EPO-H is also expected to have enhanced pharmacokinetics. Fusing mutated EPO to the IH4 nanobody not only increases the size of the molecule to avoid renal clearance, but it also directs the fusion protein to mature RBCs in circulation, further extending serum half-life (Fig. 1J) (Kontos and Hubbell, 2010; Kontos *et al*., 2013; Burrill *et al*., 2016). Through these strategies, we constructed EPO-H fusion proteins that would allow administration of high doses required for tissue protection but avoid thrombosis, thereby achieving prolonged activity in the body and reduced dosing frequency.

### Erythropoietic activity of EPO variants *in vitro*

The ability of different EPO mutants to promote RBC production was tested *in vitro* via TF-1 cell proliferation assays. TF-1 is an immature erythroid cell line that expresses both EPOR and GPA (1620 ± 140 and 3860 ± 780 molecules per cell, respectively) (Taylor *et al*., 2010), and requires EPO, GM-CSF, or IL-3 for growth (Kitamura *et al*., 1989). TF-1 cells were starved of cytokines overnight and then exposed to EPO variants for 72 hr. Their proliferation was measured by a standard tetrazolium-based assay. Wild-type EPO (epoetin alfa) and hyperglycosylated EPO (darbepoetin) exhibited EC_50_ values of ∼0.1 nM and ∼1 nM, respectively. EPO mutations on the strong binding face reduced activity of unfused EPO by ∼120- to 3400-fold relative to epoetin alfa. When these mutants were fused to IH4, their activities were rescued by ∼180- to 340-fold relative to unfused mutants, showing comparable activity to epoetin alfa and darbepoetin (Table II and Fig. S1). The unfused EPOs with mutations on the weak face showed no activity at concentrations ranging from 10^−14^ to 10^−7^ M, except for EPO(R103K), which had a slightly lower EC_50_ value compared to epoetin alfa and approximately two-fold lower efficacy (E_max_) (Table II and Fig. S1). When EPO(R103K) was fused to IH4, the fusion protein exhibited significantly enhanced activity, with its EC_50_ value in a low femtomolar range. Among the weak-face mutants that completely lacked erythropoietic activity, only IH4-EPO(L108A) exhibited targeted erythropoietic activity, while the others remained inactive even after fusion. Similar to IH4-EPO(R103K), IH4-EPO(L108A) also had an EC_50_ of ∼1–10 fM (Table II and Fig. 2A).

**Fig. 2.**
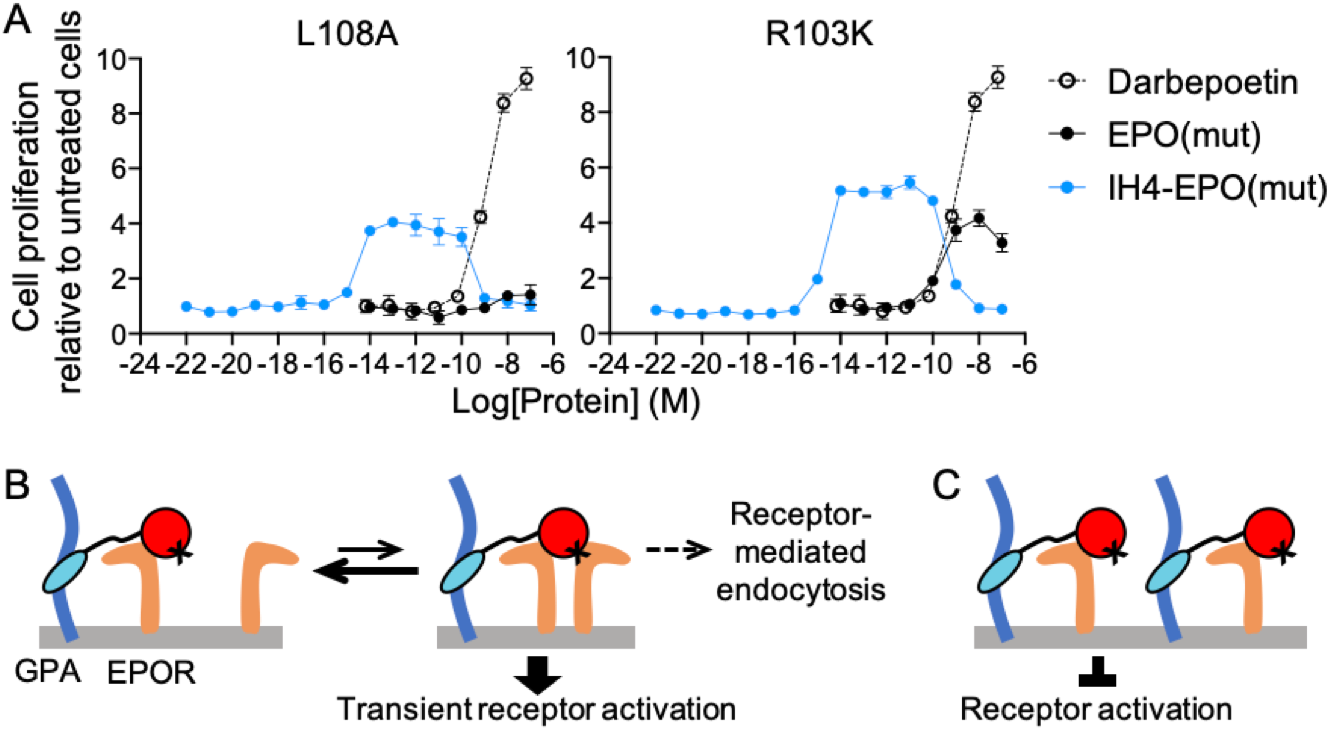
Receptor activation by IH4-EPO(L108A or R103K) in TF-1 cells follows a bell-shaped dose response curve. **(A)** IH4-EPO(L108A or R103K) was tested for stimulation of proliferation of TF-1 cells, which express both EPOR and GPA (Taylor *et al*., 2010). The fusion proteins show extremely high potency, with EC_50_ values at a low femtomolar range, and a drop in bioactivity at high concentrations. Data represent mean ± S.E.M. of three replicates. **(B**,**C)** Schematic of proposed mechanisms for the bell-shaped dose response curve. The fusion protein binds to GPA (*dark blue*) and one copy of EPOR (*orange*) via IH4 (*light blue*) and the strong face of EPO (*red*), respectively. **(B)** At low fusion protein concentrations, EPO has a brief interaction with the second copy of EPOR via the EPO weak face. This transient interaction activates EPOR signaling for cell proliferation, but does not last long enough to trigger receptor-mediated endocytosis. Thus, signaling does not terminate and a few signaling complexes per cell are sufficient to stimulate proliferation. **(C)** At high concentrations, fusion proteins saturate EPORs with a 1:1 stoichiometry via the strong-face interaction, resulting in a low number of complete signaling complexes composed of one ligand and two receptors.

The dose-response curve of weak-face mutants fused to IH4 showed two unusual features. First, when EPO(L108A or R103K) is fused to IH4, the potency of the fusion protein is enhanced by four to five orders of magnitude relative to wild-type EPO and other EPO fusion proteins. The EC_50_ is ∼1–10 fM (Fig. 2A). Secondly, the dose-response curve of IH4-EPO(L108A or R103K) is bell-shaped, with stimulation falling off at ∼1 nM, whereas fusion proteins containing strong-face mutants (K45D and R150A) show standard sigmoidal dose-response curves (Fig. 2A and Fig. S1). We speculate that these features result from distinct receptor binding properties of weak-face mutants. These mutations further reduce EPO– EPOR interaction at the weak face, resulting in an extremely rapid off-rate. At low concentrations of the fusion protein, the binding of EPOR to EPO’s weak face, needed for the formation of a complete signaling complex, may be so transient that the fusion protein activates EPORs for cell proliferation but cannot stay long enough to be endocytosed. This has the net effect of increasing the frequency of EPOR activation with a limited amount of the fusion protein (Fig. 2B). At high concentrations, the fusion protein saturates EPORs in a non-signaling, monomeric form via the strong side, and blocks receptor activation (Fig. 2C).

### Targeted erythropoietic activity of EPO-H in mice

One of the fusion proteins, IH4-EPO(L108A), was tested for targeted erythropoietic activity in transgenic mice expressing human GPA. IH4-EPO(L108A) was chosen because EPO(L108A) by itself showed essentially no homodimeric EPOR activation in TF-1 cell proliferation assay, suggesting that potential pro-thrombotic side effects would be greatly reduced. Mice received a single intraperitoneal (i.p.) injection of saline, darbepoetin (50 pmol = 2 μg; 12.5 pmol = 0.5 μg) or IH4-EPO(L108A) (6 pmol = 0.3 μg). Target cell specificity and drug efficacy were measured by staining for reticulocytes and reticulated platelets in blood samples on Days 0, 4 and 7 post-injection. Reticulocyte and reticulated platelet levels remained at baseline (Day 0) throughout the experiment in the saline-treated mice, but increased significantly in mice treated with darbepoetin, a control for the untargeted form of EPO. Mice treated with IH4-EPO(L108A) had elevated reticulocyte counts (8.47%) that were comparable to those in mice treated with 12.5 pmol of darbepoetin (8.17%) on Day 4 (Fig. 3A), but did not have significantly increased reticulated platelet counts (Fig. 3B). When various doses were tested (40 pmol = 2 μg; 6 pmol = 0.3 μg; 1.2 pmol = 0.06 μg; 0.2 pmol = 0.01 μg), IH4-EPO(L108A) induced reticulocyte responses in a dose-dependent manner: 40 pmol and 6 pmol resulted in 8.13% and 3.16% increases in reticulocyte counts on Day 4 relative to Day 0, respectively, while lower doses (1.2 pmol and 0.2 pmol) did not have significant effects (Fig. S2).

**Fig. 3.**
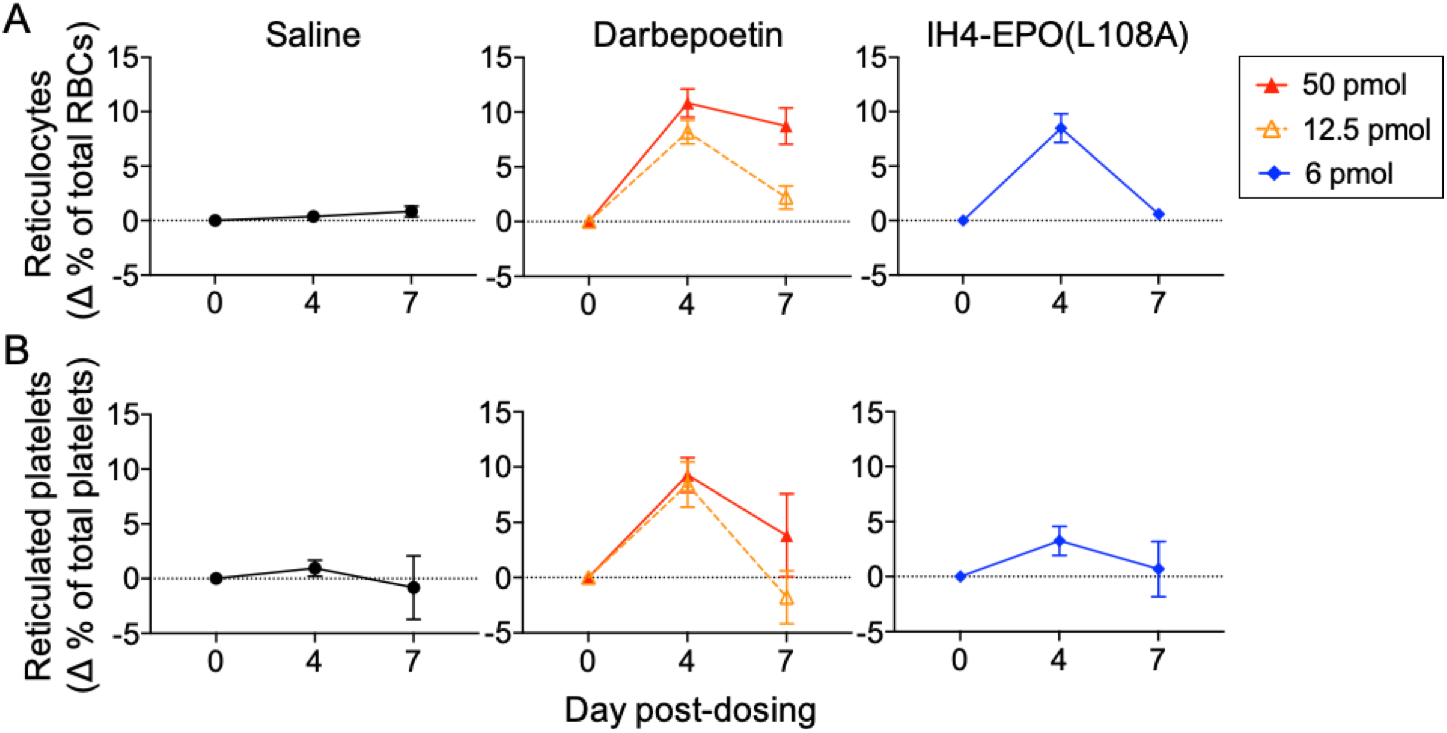
Erythropoietic activity of IH4-EPO(L108A) *in vivo*. Transgenic mice that express human GPA on their RBCs received a single i.p. injection of saline, darbepoetin or IH4-EPO(L108A). Their reticulocyte and reticulated platelet levels were measured by flow cytometry on Days 0, 4 and 7 post-injection. **(A**,**B)** While untargeted form, darbepoetin, stimulates the production of both reticulocytes and reticulated platelets, IH4-EPO(L108A) specifically stimulates RBC production and not platelet production in these mice. Data represent mean ± S.E.M of five mice per dose group.

### Tissue-protective activity of EPO variants *in vitro*

EPO mutants that displayed targeted erythropoietic activity were further evaluated to confirm their expected tissue-protective effects in cell-based assays. The ability of a fusion protein to protect cells was measured *in vitro* by estimating the number of surviving cells after treatment with EPO and a cobalt chloride (CoCl_2_), which induces a hypoxia response via hypoxia-inducible factor-alpha (HIF-1α) due to its inhibition of prolyl hydroxylase (Epstein *et al*., 2001; Vengellur and LaPres, 2004; Yuan *et al*., 2003). SH-SY5Y, a neuroblastoma cell line that expresses both EPOR and CD131 (Chamorro *et al*., 2013), was co-treated with engineered EPO variants and 100 μM of CoCl_2_. Viable cells were measured 24 hr later by standard tetrazolium dye-based assays. The optimal cell density and concentration of CoCl_2_ were chosen to cause ∼30–40% cell viability in the absence of EPO.

The control proteins, wild-type EPO (EPO(WT)) and EPO(S104I), protect neuroblastoma cells from CoCl_2_ insult, although EPO(S104I) had a much weaker effect than the wild-type (Fig. 4A, B and Fig. S3). This is consistent with the previous results that EPO(WT) and EPO(S104I) protected primary neurons from N-methyl-D-aspartic acid (NMDA)-induced excitotoxicity (Gan *et al*., 2012). While a fusion protein containing a strong-face mutant, IH4-EPO(K45D), did not protect cells from CoCl_2_-induced cell death (Fig. 4C and Fig. S3), fusion proteins containing a weak-face mutant, IH4-EPO(R103K) and IH4-EPO(L108A), exhibited neuroprotective effects (Fig. 4D, E and Fig. S3). Similarly, a weak-face mutant, EPO(L108A), also showed neuroprotective effect against CoCl_2_-induced hypoxic damage in the absence of fusion to the IH4 nanobody (Fig. 4F and Fig. S3). Four-parameter fits of these data did not give accurate EC_50_ values because the readouts did not reach saturation within the measured concentration range. Despite this caveat, four-parameter fits provided rough estimates for the potency of each variant. The EC_50_ values of EPO(WT) and EPO(S104I) were ∼1–5 nM. The EC_50_ values of IH4-EPO(R103K), IH4-EPO(L108A) and EPO(L108A) were estimated to be ∼10–20 nM (Fig. 4G).

**Fig. 4.**
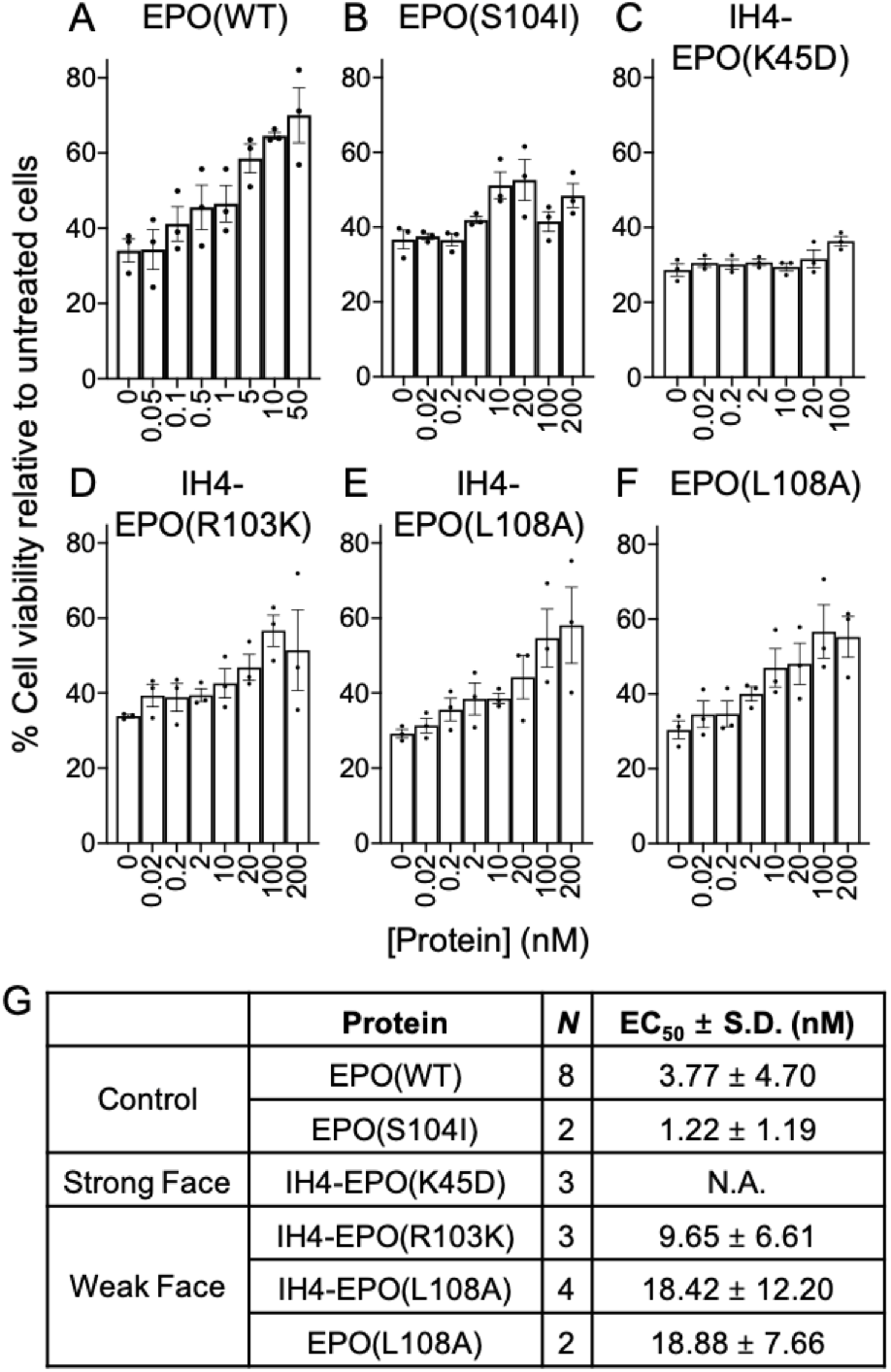
Ability of EPO variants to protect neuronal cells from CoCl_2_-induced hypoxic damage *in vitro*. SH-SY5Y cells were co-treated with EPO and CoCl_2_ for 24 hr and cell viability was measured. **(A**,**B)** Positive controls, EPO(WT) and EPO(S104I), protect neuronal cells from cell death in a dose-dependent manner, but **(C)** fusion protein containing a strong-face mutant, IH4-EPO(K45D), does not promote neuroprotection. **(D–F)** EPO variants containing a weak-face mutation, EPO(L108A) and IH4-EPO(L108A or R103K), also show neuroprotective effect against CoCl_2_-induced hypoxic damage. Data represent mean ± S.E.M. of three replicates. **(G)** A summary of tissue-protective activity of EPO variants. *N* indicates the number of repeat experiments, each containing two to four replicates. EC_50_ values were estimated by the standard four-parameter non-linear fit. See Supplementary Information for more details.

Although the dynamic range and the four-parameter fits varied between independent experiments, these EPO variants showed reproducible effects when they were repeated several times and even when they were tested under different experimental conditions (Fig. S3–S5). When SH-SY5Y cells were pre-exposed to EPO variants 24 hr before receiving 100 μM of CoCl_2_, EPO(L108A or R103K) in both unfused and fused forms protected cells from hypoxia-induced cell death (Fig. S4).

## Discussion

In this work, we constructed an EPO fusion protein that can provide both erythropoietic and tissue-protective effects without causing thrombotic side effects. This novel molecule is designed to prevent or treat hypoxia-mediated damage in patients suffering from illnesses, such as COPD and right-side heart failure, to prevent altitude sickness in military personnel acclimating to high altitude regions and possibly to enhance physical performance. It may also alleviate organ damage caused by hypoxia in COVID-19 patients at risk of requiring a ventilator. However, achieving a safe and tissue-protective dose of EPO is a challenge: the maximum allowed dose in patients with chronic kidney disease is limited by its pro-thrombotic effects (Nichol *et al*., 2015), and the dose of EPO required for tissue-protective effects is at least as high or higher than for erythropoiesis (Masuda *et al*., 1994). If doses are limited to non-thrombotic “safe” levels, then EPO is likely to fail in clinical trials for tissue protection because such doses are below what is effective for tissue protection and not necessarily because the drug itself is not effective. By using the chimeric activator design, we addressed two major challenges in using EPO activity for the treatment of hypoxia – retaining both the erythropoietic activity and tissue-protective functions of EPO while reducing or eliminating its pro-thrombotic activity.

Previously described EPO derivatives lack the desired features for treatment of hypoxia. Darbepoetin simply extends the plasma half-life but has the same activities as EPO itself (Egrie and Browne, 2001; Egrie *et al*., 2003). Carbamylated EPO (Leist *et al*., 2004) and weak-face EPO mutants (such as EPO(S104I) (Gan *et al*., 2012)) retain neuroprotective activity but completely lack erythropoietic and pro-thrombotic activity. Targeted EPO molecules that we have constructed (Burrill *et al*., 2016; Lee *et al*., 2020) retain erythropoietic activity and are not pro-thrombotic, but are predicted to lack tissue-protective activity because they contain a mutation in the surface of EPO that strongly binds to EPOR; this surface is predicted to be critical for binding to EPOR–CD131 heterodimers that mediate tissue protection.

We therefore designed new EPO derivatives by combining two features: (1) a mutation on the surface of EPO that interacts weakly with EPOR, since such mutant EPOs retain tissue-protective activity and (2) an antibody-based GPA-binding element that should rescue activity on RBC precursors. This design mimics our previous Targeted EPO, except that the EPO mutation is in the weak face instead of the strong face with respect to interaction with EPOR. We found that most mutations in the weak face (specifically R14E, R14Q, R14N, Y15I, R103I and R103Q) abolished erythropoietic activity in cell-based proliferation assays, and the mutation R103K caused only a slight reduction in this assay. In contrast, EPO containing the mutation L108A completely lacked *in vitro* erythropoietic activity, but this activity was rescued when the mutant protein was fused to the GPA-targeting nanobody IH4 (Table II, Fig. 2A and Fig. S1). In addition, this fusion protein retained erythropoietic activity *in vivo* (Fig. 3 and Fig. S2), with a potency similar to our Targeted EPO and EPO itself (Burrill *et al*., 2016; Lee *et al*., 2020). Finally, the IH4-EPO(L108A) fusion protein still showed neuroprotective activity *in vitro* in an assay in which neuroblastoma cells were treated with CoCl_2_, which induces a hypoxia response (Fig. 4, Fig. S3 and Fig. S4). Thus, the fusion protein IH4-EPO(L108A) is a potential candidate for treatment of hypoxia.

In *in vitro* testing for erythropoietic activity, the IH4-EPO(L108A) and IH4-EPO(R103K) fusion proteins showed two unusual properties: (1) extreme potency, with activity detectable at ∼1–10 fM concentrations, and (2) loss of activity at ∼1 nM concentration (Fig. 2A). These observations were made based on TF-1 erythroleukemia cell proliferation assays, in which cells were stimulated by wild-type EPO and engineered proteins. These effects are likely not relevant *in vivo*, since IH4-EPO(L108A) stimulates erythropoiesis at doses similar to EPO itself and does not show signs of extreme potency or auto-inhibition (Fig. 3 and Fig. S2). Nonetheless, it may be useful to have a working model of these observations, as such an understanding may inform the engineering of other targeted proteins.

We propose two mechanisms that could, in combination, explain the extremely high potency of IH4-EPO(L108A or R103K). First, the attachment of the fusion protein to GPA could prevent receptor-mediated endocytosis and degradation of the signaling protein. In differentiating erythroleukemic cells, GPA is attached to a stable actin cytoskeleton that may preclude internalization. In non-differentiating erythroleukemic cells, GPA is internalized by a clathrin-mediated pathway but at a much slower rate compared to other membrane proteins (Marshall *et al*., 1984; Ktistakis *et al*., 1990). Therefore, binding to GPA may interfere with internalization and degradation of the fusion protein. However, simple attachment of an EPO fusion protein to GPA does not profoundly enhance its potency, since we do not observe highly potent activity with other anti-GPA/EPO fusion proteins. Second, we propose that the fusion protein may form a highly stable complex with GPA and one copy of EPOR via the strongly interacting side of EPO, but interaction with the second EPOR to form a complete signaling complex may be weak and dissociate rapidly due to a mutation. The interaction may last long enough to phosphorylate a subset of the tyrosine residues important in signal transduction into the nucleus but may not be long enough to phosphorylate residues involved in signaling to the clathrin system for receptor-mediated endocytosis (Fig. 2B). In these assays, stimulation of proliferation is observed with as few as six to sixty molecules of fusion protein per cell, suggesting that this is the minimum number of molecules needed to promote erythropoietic signaling (see Supplementary Information for a quantitative explanation).

The loss of activity by IH4-EPO(L108A or R103K) at >1 nM concentrations can be explained by receptor saturation that has been observed in other systems that require more than two receptors for signaling (Fuh *et al*., 1992; Atanasova and Whitty, 2012; Kallenberger *et al*., 2014). Similar to EPO, human growth hormone (hGH) asymmetrically binds to two hGH receptors to trigger signaling. Fuh *et al*. (1992) showed that wild-type hGH inhibits signaling at >2 μM, and mutation of the weak-binding face of hGH further reduces the IC_50_ value to ∼100 nM. They also demonstrated that the antagonistic behaviors resulted from the disruption of receptor dimerization, using divalent monoclonal antibodies for hGH receptors (Fuh *et al*., 1992). Similarly, our fusion protein containing a weak-face EPO mutation may saturate monomeric EPOR in a 1:1 stoichiometry and block the formation of a complete signaling complex consisting of homodimeric EPOR, resulting in auto-inhibition of EPO signaling (Fig. 2C).

It is important to note that while we observe an enhanced potency of IH4-EPO(L108A) in the cell based assay, in our *in vivo* erythropoiesis experiments the potency of this molecule was similar to that of darbepoetin and EPO fusion proteins that we constructed previously. In the *in vitro* assay, there may be essentially no removal of the fusion protein, while *in vivo* the normal clearance mechanisms would likely still operate, such as pinocytosis and degradation by Kupffer cells and/or binding to EPORs on non-erythroid cells and removal by non-signaling receptor-mediated endocytosis (Wiley, 2003).

Taken together, our results indicate that IH4-EPO(L108A) could be an ideal molecule for treatment of hypoxia. The fusion protein is expected to enhance oxygen delivery and prevent hypoxia-induced cell death, without causing thrombosis. This work demonstrates that our engineering strategies allow for selective utilization of beneficial EPO activities and inhibition of undesired effects. More broadly, it further solidifies the value of the “chimeric activator” approach in designing targeted protein therapeutics.

## Methods

### Cell culture

FreeStyle 293-F and FreeStyle CHO-S cell lines were obtained from Invitrogen (Carlsbad, CA) and cultured in FreeStyle 293 Expression Medium and complete FreeStyle CHO Expression Medium (Invitrogen), respectively. Human erythroleukemia TF-1 and human neuroblastoma SH-SY5Y were obtained by ATCC (Manassas, VA). TF-1 was cultured in RPMI-1640 with 10% FBS, 100 U/mL penicillin, 100 U/mL streptomycin, and 2 ng/mL recombinant human granulocyte macrophage colony-stimulating factor (GM-CSF; PeproTech) unless specified otherwise. SH-SY5Y was cultured in 1:1 DMEM/F-12 with 10% FBS. 293-F and CHO-S were cultured at 37°C in 8% CO_2_ with shaking at 2.35 x g. TF-1 and SH-SY5Y were cultured at 37°C in 5% CO_2_.

### DNA constructs

The DNA sequence for EPO wild-type was from GenBank (accession no. KX026660). EPO mutant sequences were constructed by introducing a codon change into the wild-type sequence. The DNA sequence for the IH4 nanobody was derived by reverse translating and codon optimizing (Integrated DNA Technologies) the protein sequence adapted from the US patent 9879090 (Bertrand *et al*., 2018). It was modified to include a point mutation (Phe80Tyr) in the framework region 3 to reflect the consensus of the germline sequences, and an additional amino acid (Thr118) in the framework region 4, as the reported sequence had a typographical error. See Supplementary Information for individual sequences.

### Protein expression and purification

Transient expression was performed in 293-F and CHO-S cells using pSecTag2A or pOptiVEC plasmids according to the supplier’s protocol. 4–6 days after transfection, protein expression was assayed by Western blotting cell supernatant using anti-6xHis-HRP antibody (Abcam). Proteins from transient transfection were purified as follows. Supernatant was concentrated to 5–8 mL using a 10 kDa cut-off Macrosep Advance centrifugal device (Pall). Concentrated protein was bound to 0.5–1 mL of His60 nickel or HisTalon cobalt resin (Takara Bio) for 0.5–1 hr at 4°C while rotating in a 10-mL Pierce disposable column (Thermo Scientific), and was washed and eluted using His60 or HisTalon Buffer Set (Takara Bio) according to the supplier’s protocol. Cell supernatant and each purification fraction were analyzed by SDS–PAGE followed by Coomassie Blue staining. Eluted proteins were combined, desalted into endotoxin-free PBS (Teknova: 137 mM NaCl, 1.4 mM KH_2_PO_4_, 4.3 mM Na_2_HPO_4_, and 2.7 mM KCl, pH 7.4) using Econo-Pac 10DG columns (Bio-Rad), and concentrated to <1 mL using Macrosep Advance centrifugal device.

For *in vivo* experiments, contaminating proteins were further removed by anion-exchange chromatography (AIEX) on HiPrep DEAE FF 16/10, followed by size exclusion chromatography (SEC) on Superdex 200 10/300 GL columns (Cytiva), using AKTA FPLC system (Cytiva). For AIEX, 1 M Tris-HCl, pH 8.0 was used as the starting buffer and a linear gradient up to 1 M NaCl was used for elution. For SEC, endotoxin-free PBS was used as the running buffer. Desired protein fractions were combined and concentrated to <1 mL using Macrosep Advance centrifugal device. Proteins were stored at 4°C throughout the described process, ultimately stored as aliquots at −80°C, and thawed once before use. Only endotoxin-free reagents were used.

### TF-1 cell proliferation assays

TF-1 cells were seeded in a 96-well plate at 9.0×10^3^ cells per well in 90 μL of RPMI-1640 with serum and antibiotics (no GM-CSF). The purified proteins were serially diluted by 10-fold (10^−7^ to 10^−14^ or 10^−21^ M) or 100-fold (10^−7^ to 10^−21^ M) and added to the cells. Cells were incubated at 37°C in 5% CO_2_ for 72 hr. Cell proliferation was determined by CellTiter 96^®^ AQueous One Solution Cell Proliferation Assay (Promega) or adding 10 μL of WST-1 reagent (Roche). 2–4 hr after adding the reagent, absorbance at 490 nm (and background absorbance at 650 nm when using WST-1) was read on a BioTek Synergy Neo HTS microplate reader. Reported data represent mean ± SEM of three replicates.

### Measuring mouse reticulocytes and reticulated platelets

Human GPA-transgenic FVB mice were generously donated by the Hendrickson Laboratory at Yale University (Auffray *et al*., 2001). This strain underwent embryo re-derivation at Charles River Laboratories. The homozygous human GPA transgene is embryonic-lethal but heterozygotes are phenotypically normal, so a breeding colony was maintained with screening for human GPA at each generation. Transgene expression was measured as described before (Burrill *et al*., 2016).

Five mice per dose group received a single intraperitoneal (i.p.) injection with saline, darbepoetin or EPO fusion protein in a 200 μL volume (diluted in saline or PBS) on Day 0. 1–5 μL of whole blood was collected by tail-nick in EDTA-coated tubes on Days 0, 4 and 7 post-injection. Blood was analyzed immediately after collection by flow cytometry as described before (Burrill *et al*., 2016). Briefly, thiazole orange (Sigma-Aldrich) was used to stain residual RNA in reticulocytes and reticulated platelets, and anti-CD41-PE antibody (BD Pharmingen) was used to stain total platelets. A stock solution (1 mg/mL) of thiazole orange was prepared in 100% methanol and was diluted 1:5,000 in PBS to make a 2x working solution. Anti-CD41-PE antibody was diluted 1:500 in either the 2x working solution of thiazole orange for stained samples or PBS for gating thiazole orange-negative population. Whole blood was diluted 1:1,000 in PBS. Equal volumes (100 μL) of 2x working solution of anti-CD41-PE antibody with or without thiazole orange and diluted whole blood were mixed in a 96-well U-bottom plate and incubated for 30 min in the dark at 23 °C. The fluorescence was measured on a LSRFortessa SORP flow cytometer equipped with an optional HTS sampler (BD Biosciences) using the following filter configuration: PE excitation, 561/50 mW; emission filter, BP 582/15; YFP excitation, 488/100 mW; emission filter, BP 540/25.

### Tissue protection assay

SH-SY5Y cells were seeded in a 96-well plate at 4.8×10^4^ cells per well in 80 μL of 1:1 DMEM/F-12 with 10% FBS, and let adhere overnight at 37°C in 5% CO_2_. In co-treatment experiments, cells received varying concentrations of purified proteins (0.02 to 200 nM) and 100 μM of cobalt chloride (CoCl_2_), a hypoxia mimicking agent, and were incubated at 37°C in 5% CO_2_ for 24 hr. In pre-treatment experiments (Fig. S4), cells were treated with purified proteins 24 hr before receiving CoCl_2_ and were incubated at 37°C in 5% CO_2_ for additional 24 hr after adding CoCl_2_. Cell viability was measured by CellTiter 96® AQ_ueous_ One Solution Cell Proliferation Assay (Promega). 2–4 hr after adding the reagent, absorbance at 490 nm was read on a BioTek Synergy Neo HTS microplate reader. In experiments shown in Supplementary Figure 5, SH-SY5Y cells were seeded in a 96-well plate at 1.2×10^4^ cells per well in 80 μL. On the next day, cells were co-treated with varying concentrations of purified proteins (0.02 to 200 nM) and 25 or 50 μM of CoCl_2_, and incubated at 37°C in 5% CO_2_ for 72 hr. Cell viability was measured by adding 10 μL of WST-1 reagent (Roche). 4 hr after adding the reagent, absorbance at 490 nm and 650 nm (background) was read on a BioTek Synergy Neo HTS microplate reader. Reported data represent mean ± S.E.M of two to four replicates.

## Acknowledgements

This work was supported by funds from the Wyss Institute for Biologically Inspired Engineering and the Boston Biomedical Innovation Center (Pilot Award 112475; Drive Award U54HL119145). J.L., K.M.K., D.R.B., J.C.W. and P.A.S. were supported by the Harvard Medical School Department of Systems Biology. J.C.W. was further supported by the Harvard Medical School Laboratory of Systems Pharmacology. A.V., D.R.B. and P.A.S. were further supported by the Wyss Institute for Biologically Inspired Engineering. N.G.G. was sponsored by the Army Research Office under Grant Number W911NF-17-2-0092. The views and conclusions contained in this document are those of the authors and should not be interpreted as representing the official policies, either expressed or implied, of the Army Research Office or the U.S. Government. The U.S. Government is authorized to reproduce and distribute reprints for Government purposes notwithstanding any copyright notation herein. We sincerely thank Amanda Graveline and the Wyss Institute at Harvard for their scientific support.

## Author contributions

J.L., K.M.K. and J.C.W. designed research; J.L., A.V., N.G.G., K.M.K. and J.C.W. performed research;

J.L., K.M.K., D.R.B. and J.C.W. analyzed data; and, J.L., D.R.B., J.C.W. and P.A.S. wrote the paper.

## Conflict of Interest Statement

J.C.W., D.R.B. and P.A.S. are shareholders in a company that has a license to the IH4 antibody element described in this work.

