## Supplementary Information for "Rational engineering of an erythropoietin fusion protein to treat hypoxia"

**Table of Contents**

Supplementary Figures (Fig. S1–5)
Quantitative explanation of extreme potency of IH4-EPO(L108A or R103K)
Table of Relevant Binding Parameters
DNA and Protein Sequences
EPO Mutations

**Supplementary Figures**

Supplementary Figure 1

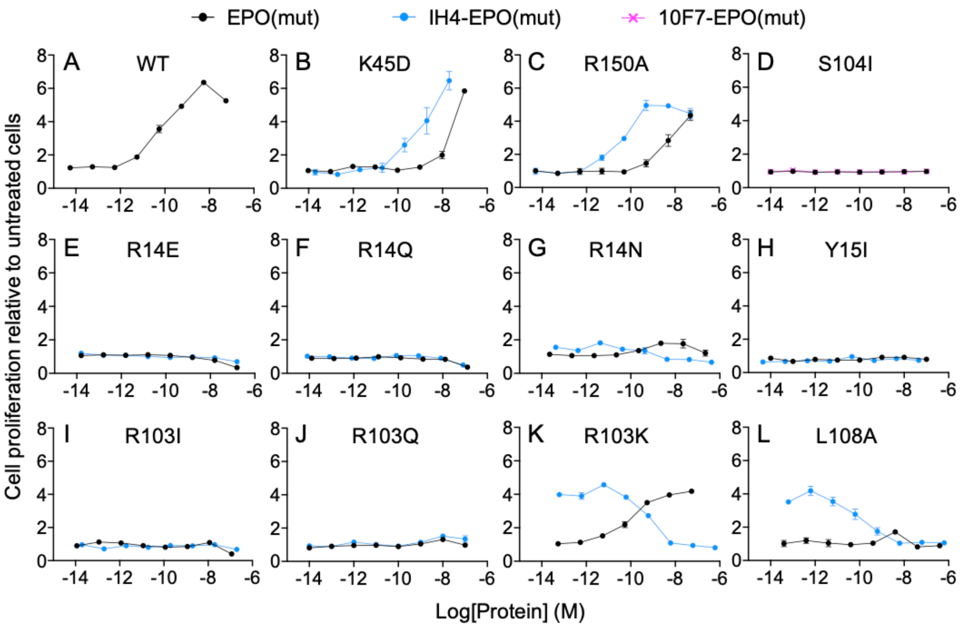


**Fig. S1.** *In vitro* erythropoietic activities of EPO mutants in unfused and antibody-fused forms. Standard TF-1 cell proliferation assays were performed to measure the ability of EPO mutants to stimulate cell proliferation. The same mutants fused to an anti-GPA antibody fragment (IH4 nanobody or 10F7 scFv) via a five-amino acid linker were tested for GPA-dependent activation of EPOR. **(A)** Wildtype EPO activity as a positive control. **(B,C)** Strong-face mutants, K45D and R150A, reduce the EPO activity but show enhanced activity upon fusion to IH4. **(D–J)** Most weak-face mutations at R14, Y15, S104, and R103 of EPO do not show any activity even when they are fused to an antibody element. **(K,L)** Weak-face mutants, R103K and L108A, show slightly reduced and almost no activity by itself, respectively. Their fusion to IH4 exhibits inverted dose response curves, in which their activity is greatly enhanced at low concentrations but drops back to the baseline at high concentrations. Data represent mean ± S.E.M. of three replicates.

Supplementary Figure 2

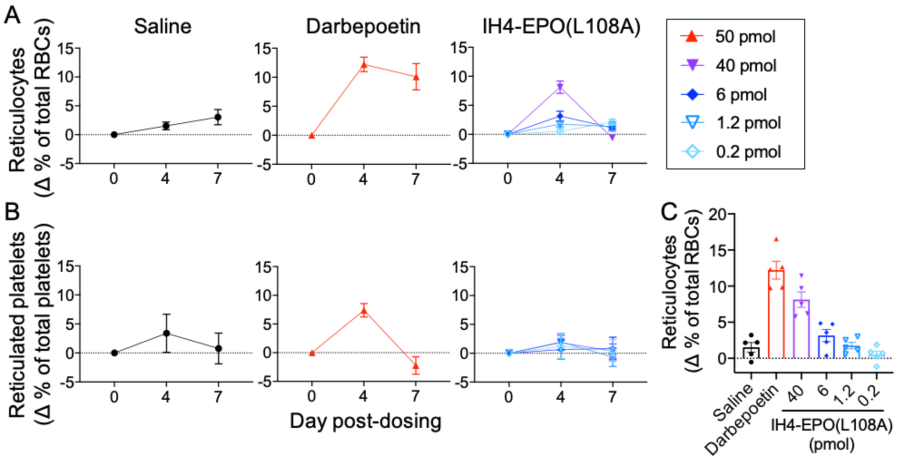


**Fig. S2.** *In vivo* erythropoietic activity of IH4-EPO(L108A). Transgenic mice that express human GPA on RBCs received a single i.p. injection of saline, darbepoetin, or IH4-EPO(L108A). Reticulocyte and reticulated platelet levels were measured by flow cytometry on Days 0, 4, and 7 post-injection. **(A,B)** IH4-EPO(L108A) specifically stimulates RBC production and not platelet production. **(C)** IH4-EPO(L108A) induces erythropoiesis in a dose-dependent manner as shown on Day 4 post-injection. Data represent mean ± S.E.M of five mice per dose group.

Supplementary Figure 3

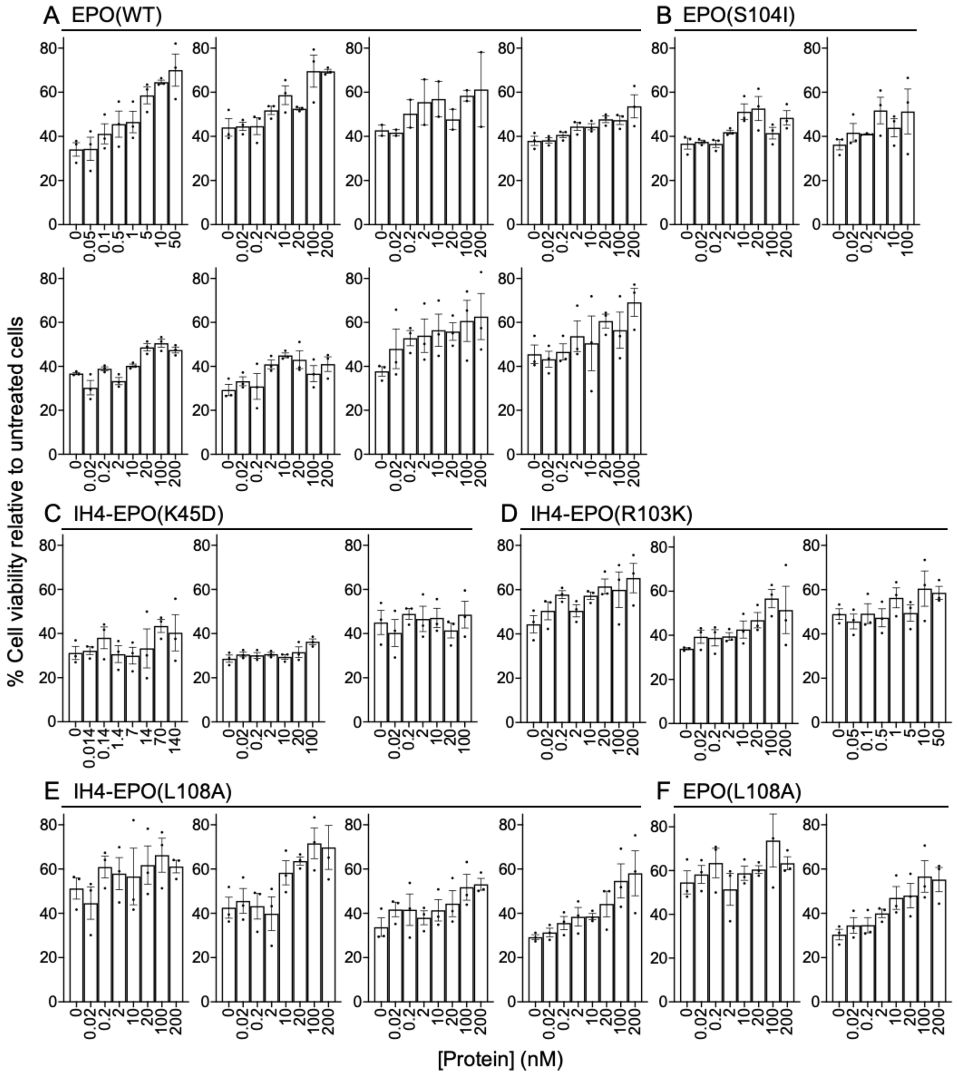


**Fig. S3.** Ability of EPO variants to protect neuronal cells from $CoCl_2$-induced hypoxic damage *in vitro*. SH-SY5Y cells were co-treated with EPO and 100 µM $CoCl_2$ for 24 hr and cell viability was measured. For each protein, at least two repeat experiments were performed. **(A,B)** Two positive controls, EPO(WT) and EPO(S104I), show tissue-protective effect in a dose-dependent manner, although EPO(S104I) shows much weaker effect. **(C)** Fusion protein containing a strong-face mutation, K45D, does not have tissue-protective activity. **(D,E)** Fusion protein containing a weak-face mutation, R103K or L108A, protects neuronal cells from $CoCl_2$-induced cell death. Note that the dynamic range can vary between experiments but the tissue-protective effects are reproducible. Data represent mean ± S.E.M. of two to three replicates.

Supplementary Figure 4

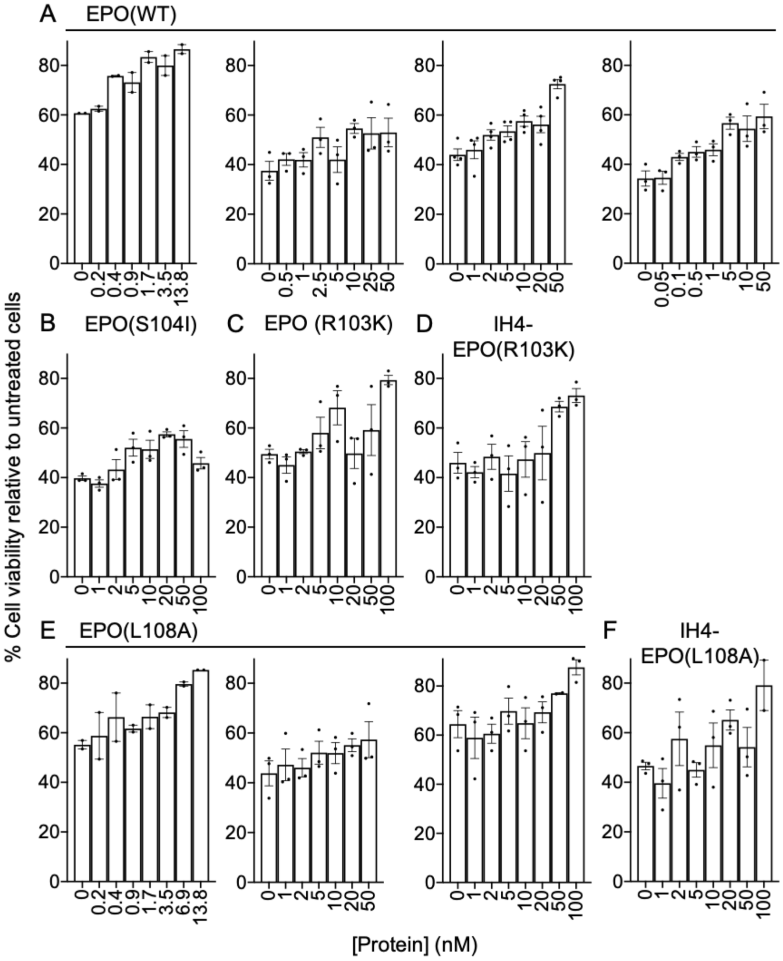


**Fig. S4.** Pre-exposure to EPO variants also protects neuronal cells from $CoCl_2$-induced hypoxic damage *in vitro*. SH-SY5Y cells were pre-treated with EPO 24 hr prior to adding $CoCl_2$. Cells were incubated for additional 24 hr after receiving 100 μM $CoCl_2$ and cell viability was measured. **(A,B)** Two positive controls, EPO(WT) and EPO(S104I), show tissue-protective effect in a dose-dependent manner, although EPO(S104I) shows much weaker effect. **(C–F)** EPO(R103K or L108A) in an unfused or antibody-fused form protects neuronal cells from $CoCl_2$-induced cell death. Note that the dynamic range can vary between experiments but the tissue-protective effects are reproducible. Data represent mean ± S.E.M. of two to four replicates.

Supplementary Figure 5

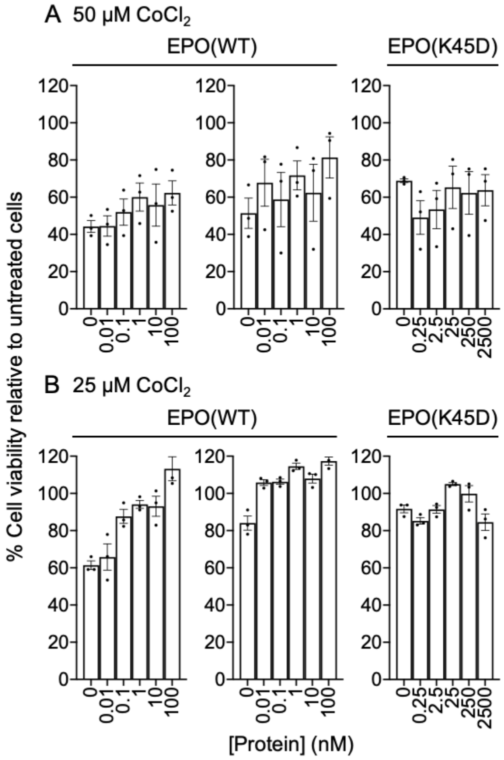


**Fig. S5.** Ability of EPO variants to protect SH-SY5Y cells from $CoCl_2$-induced hypoxic damage *in vitro*. SH-SY5Y cells were co-treated with EPO and $CoCl_2$ for 72 hr and cell viability was measured. Hypoxic damage was induced using **(A)** 50 μM and **(B)** 25 μM $CoCl_2$. Positive control, EPO(WT), but not a strong-side mutant, EPO(K45D), shows tissue-protective activity. Note that the dynamic range can vary between experiments but the tissue-protective effects are reproducible. Data represent mean ± S.E.M. of three replicates.

**Quantitative explanation of extreme potency of IH4-EPO(L108A or R103K)**

We hypothesize that the extremely potent signaling of IH4-EPO(L108A or R103K) (Table II and Fig. 2A) can be explained by a lack of receptor-mediated endocytosis, such that signaling is not terminated after receptor activation (Fig. 2B). Some receptor tyrosine kinase systems involve a rapid phosphorylation event(s) that initiates signaling that results in transcriptional modulation, followed by slower phosphorylation events that lead to receptor-mediated endocytosis and degradation of the receptor and/or ligand (Wiley, 2003). According to our hypothesis, mutation of the weak face of EPO could further reduce the stability of the EPO–$(EPOR)_2$ complex, such that this complex is rapidly forming but also dissociates so rapidly that endocytosis-stimulating phosphorylation does not occur.

Despite the extreme potency of IH4-EPO(L108A) *in vitro*, this fusion protein does not have enhanced potency *in vivo*, and behaves similarly to our previous Targeted EPO (Burrill *et al*., 2016; Lee *et al*., 2020) with respect to RBC production and lack of platelet production (Fig. 3). In a culture dish, receptor-mediated endocytosis by a single cell type is the only mechanism to terminate signaling, but *in vivo* the fusion protein may disappear through renal clearance, bulk fluid pinocytosis, and by binding to EPOR on non-hematopoietic cells, followed by non-signaling endocytosis and protein degradation that is part of normal membrane turnover. Thus, IH4-EPO(L108A) should have a clearance rate and potency that would be compatible with its use as a treatment for hypoxia and related disorders.

The following calculations provide quantitative explanations for our hypothesis. Kinetics parameters relevant to these calculations can be found in the “Table of Relevant Binding Parameters” below.

**1. Minimum number of fusion protein molecules needed for signaling**

The extreme potency of IH4-EPO(L108A or R103K) allows an estimate of about 6 to 60 fusion protein molecules per cell are required for EPO-induced signaling in TF-1 cells. At the $EC_{50}$ of ~1–10 fM for stimulation of TF-1 cell proliferation, there are about 6 to 60 molecules of the fusion protein per cell at the start of the assay. Specifically, there are about 9,000 cells and 600,000 fusion protein molecules per 100 μL in a well at the start of the proliferation assay. This provides an estimate for the minimum number of receptor–ligand complexes required to trigger EPO-induced signaling in TF-1 cells.

**2. Slow dissociation of a fusion protein from GPA and EPOR**

When IH4-EPO(L108A or R103K) binds to the surface of a TF-1 cell, binding is stabilized through simultaneous interaction with GPA ($K_D$ = 33 nM) and EPOR via the strong-binding face of EPO ($K_D$ = 0.1–1 nM). In this state, the local concentration of the bound fusion protein can be estimated by the number of receptor-bound fusion protein molecules and effective volume occupied by these molecules around the cell surface. The effective volume is approximated as 1 $\mu m^3$ by multiplying the cell surface area (1000 $\mu m^2$) by the distance from the cell surface in which the fusion protein is trapped (about 1 nm), so one molecule has a concentration of about $1.66 \times 10^{-9}$ M. This means that the local concentration of GPA on the surface of a TF-1 cell is about $(1.6 \times 10^{-9}$ M$) \times$ (3860 GPA molecules) = $6.2 \times 10^{-6}$ M. Given the $K_D$ value of IH4 to GPA is 33 nM, about 99.5% of GPA molecules will be also bound by fusion proteins via

IH4, when they are bound to EPOR via the strong face of the EPO element. Due to this avidity effect, the effective dissociation half-time may be increased by ~200-fold. Based on these calculations, we estimate that the effective dissociation half-time of the fusion protein from both EPOR and GPA would be ~100 hr, which is longer than the length of the experiment (72 hr). Non-signaling membrane proteins are normally endocytosed more slowly and recycling more efficiently, such that they have a metabolic turnover in the range of 12 hours or more (Wiley, 2003). Moreover, it is possible that GPA is anchored to the cytoskeleton in a way that slows or prevents this process even further (Marshall *et al*., 1984; Ktistakis *et al*., 1990). Thus, in proliferation assay wells with EPO in concentrations 1 to 100 fM, EPO will be in molar excess relative to EPO receptors, essentially all of the EPO will be bound to at least one receptor, and turnover of EPO is likely to be slow enough that much of it survives the 72-hour incubation of the assay.

#### 3. Rapid association and dissociation between a second EPOR and the GPA–EPOR–fusion protein complex

In the configuration where the fusion protein is bound to GPA and an EPOR, binding to a second EPOR would occur rapidly because the EPO element is positioned at the correct height from the membrane and in the correct orientation for such binding, and the binding would rely predominantly on the two-dimensional diffusion within the membrane. The on-rate of a fusion protein already bound to GPA and an EPOR for a second EPOR is assumed to be high because the effective molarity of the cell-bound fusion protein is high, and because the EPO element is rotationally constrained to place its weak EPOR-binding face in the correct orientation relative to the second EPOR.

However, the mutation (e.g. L108A) on the weak face of EPO likely allows for rapid dissociation of this second EPOR. The interaction with EPOR of wild-type EPO through its weak face is estimated to have a $K_D$ of about 2 μM (for the soluble interaction; Philo *et al*., 1996). Assuming that the diffusion-limited $k_{on}$ of EPO to EPOR via the weak face is the same as for the strong-face interaction ($k_{on}$ = 8.3x10$^6$ M$^{-1}$s$^{-1}$) (Gross and Lodish, 2006), the dissociation rate constant ($k_{off}$) would be about 18 s$^{-1}$, such that the complex dissociates in <0.1 second in the absence of other interactions, such as the EPOR Box1-Box2/JAK2 FERM dimerization (Ferrao *et al*., 2018).

**Table of Relevant Binding Parameters**

| Receptor | Ligand | $K_D$ (M) | $k_{on}$ ($M^{-1}s^{-1}$) | $k_{off}$ ($s^{-1}$) | $T_{1/2}$* | References |
|---|---|---|---|---|---|---|
| GPA | IH4 | $3.37x10^{-8}$ | $5.73x10^{5}$ | $1.9x10^{-2}$ | 0.6 min | Habib *et al*., 2013 |
| Soluble EPOR | EPO(WT) | $2.0x10^{-10}$ (Strong face) | – | – | – | Philo *et al*., 1996 |
| | | $2.1x10^{-6}$ (Weak face) | – | – | – | Philo *et al*., 1996 |
| EPOR on cells | EPO(WT) | $6.0x10^{-11}$ | $8.33x10^{6}$ | $5.0x10^{-4}$ | 23 min | Gross & Lodish, 2006 |
| EPOR-Fc | EPO(WT) | $5.4x10^{-9}$ | $3.9x10^{4}$ | $2.1x10^{-4}$ | 55 min | Burrill *et al*., 2016 |
| Soluble EPOR | EPO(WT) | $2.1x10^{-6}$ (Weak face) | $\sim10^{6}$ | $\sim2.1x10^{-0}$ | 0.5 sec | Inferred from Philo *et al*., 1996 |
| EPOR on cells | EPO(WT) | $6.0x10^{-7}$ (Weak face) | $\sim8.33x10^{6}$ | $\sim5x10^{0}$ | 0.2 sec | Inferred from Gross & Lodish, 2006 |
| EPOR-Fc | EPO(WT) | $5.4x10^{-5}$ | $\sim3.9x10^{4}$ | $\sim2.1x10^{0}$ | 0.5 sec | Inferred from Burrill *et al*., 2016 |
| EPOR (via weak face) | Weak-face EPO mutant | **$6.0x10^{-6}$ (Weak face) | [#]$8.33x10^{6}$ | $\sim5x10^{1}$ | 0.02 s | Estimated from Gross & Lodish, 2006 |
| EPOR (via weak face) | Weak-face EPO mutant | **$5.4x10^{-4}$ (Weak face) | [##]$3.9x10^{4}$ | $\sim2.1x10^{1}$ | 0.05 s | Estimated from Burrill *et al*., 2016 |

*For comparison, we note that the internalization rate constant ($k_{int}$) of EPO(WT) is ~$1.0x10^{-3}$ $s^{-1}$, giving an internalization half-time of ~11.5 min (Gross & Lodish, 2006) for receptor-mediated endocytosis.

**$K_D$ of the weak-face EPO L108A mutant protein to EPOR via the weak side interaction is estimated to be about 10-fold weaker than for a wild-type weak-face interaction. This is based on cell-based assay results of Elliott *et al*. (1997) and typical effects of a leucine-to-alanine mutation that removes a protein-protein interaction contact without otherwise affecting protein structure (Piehler *et al*., 2000).

[#]Diffusion-limited $k_{on}$ of a weak-face EPO mutant to EPOR is assumed to stay the same as the strong-side interaction, and is estimated based on Gross & Lodish (2006).

[##]Diffusion-limited $k_{on}$ of a weak-face EPO mutant to EPOR is assumed to stay the same as the strong-side interaction, and is estimated based on Burrill *et al*. (2016).

### DNA and Protein Sequences

| | Protein sequence | DNA sequence |
|---|---|---|
| IH4 | QVQLQESGGGSVQAGGSLRL<br>SCVASGYTDSTYCVGWFRQA<br>PGKEREGVARINTISGRPWY<br>ADSVKGRFTISQDNSKNTVY<br>LQMNSLKPEDTAIYYCTLTT<br>ANSRGFCSGGYNYKGQGTQV<br>TVS | CAGGTCCAACTGCAAGAGAGCGGCGGGGGGTCAGTTCAGGCGGGG<br>GGGAGTCTGCGGTTGAGCTGCGTAGCTTCAGGCTACACTGACAGC<br>ACCTACTGCGTGGGATGGTTTCGGCAGGCACCCGGCAAGGAACGA<br>GAGGGCGTTGCACGGATCAACACTATCTCCGGTCGGCCTTGGTAC<br>GCAGATAGTGTTAAGGGACGGTTTACTATTAGTCAGGATAACTCT<br>AAGAATACCGTCTACCTTCAGATGAATAGCCTGAAACCGGAAGAC<br>ACGGCTATTTACTATTGCACCCTTACAACTGCCAACAGCAGAGGG<br>TTTTGTTCTGGGGGATATAACTACAAAGGACAGGGGACCCAAGTC<br>ACTGTCAGC |
| 5 AA linker | SGGGS | TCTGGTGGTGGTTCC |
| EPO(WT) | APPRLICDSRVLERYLLEAK<br>EAENITTGCAEHCSLNENIT<br>VPDTKVNFYAWKRMEVGQQA<br>VEVWQGLALLSEAVLRGQAL<br>LVNSSQPWEPLQLHVDKAVS<br>GLRSLTTLLRALGAQKEAIS<br>PPDAASAAPLRTITADTFRK<br>LFRVYSNFLRGKLKLYTGEA<br>CRTGDR | GCCCCACCTAGATTGATTTGTGATTCCAGAGTTTTGGAAAGATAC<br>TTGTTGGAAGCTAAGGAGGCTGAAAATATTACTACTGGTTGTGCT<br>GAACATTGTTCTTTGAACGAGAATATTACTGTTCCAGATACTAAG<br>GTTAACTTTTACGCTTGGAAGAGAATGGAAGTTGGTCAGCAAGCT<br>GTTGAAGTTTGGCAAGGTTTGGCTTTGTTGTCTGAAGCTGTTTTG<br>AGAGGTCAAGCTTTGTTGGTTAATTCTTCTCAACCATGGGAACCA<br>TTGCAATTGCATGTTGATAAGGCTGTTTCTGGTTTGAGATCTTTG<br>ACTACCTTGTTGAGAGCTTTGGGTGCTCAAAAGGAAGCTATTTCT<br>CCTCCAGATGCTGCTTCTGCCGCTCCATTGAGAACTATTACTGCT<br>GATACTTTTAGAAAGTTGTTTAGAGTTTACTCTAACTTCTTGAGA<br>GGTAAGTTGAAGTTGTACACTGGTGAAGCTTGTAGAACTGGTGAT<br>CGG |

### EPO Mutations

| | Amino acid change | Codon change |
|---|---|---|
| K45D | Lys → Asp | AAG → GAT |
| R150A | Arg → Ala | AGA → GCC |
| R14E | Arg → Glu | AGA → GAA |
| R14Q | Arg → Gln | AGA → CAA |
| R14N | Arg → Asn | AGA → AAC |
| Y15I | Tyr → Ile | TAC → ATC |
| R103I | Arg → Ile | AGA → ATC |
| R103Q | Arg → Gln | AGA → CAG |
| R103K | Arg → Lys | AGA → AAA |
| S104I | Ser → Ile | TCT → ATC |
| L108A | Leu → Ala | TTG → GCG |
